# Role of hypocretin in the medial preoptic area in the regulation of sleep, maternal behavior and body temperature of lactating rats

**DOI:** 10.1101/2021.05.06.442961

**Authors:** Mayda Rivas, Diego Serantes, Florencia Peña, Joaquín González, Annabel Ferreira, Pablo Torterolo, Luciana Benedetto

**Author notes:** Please address correspondence to: Dr. Luciana Benedetto, Departamento de Fisiología, Facultad de Medicina, Universidad de la República, General Flores 2125, 11800 Montevideo, Uruguay.

## Abstract

The hypocretins (HCRT), also known as orexin, includes two neuroexcitatory peptides, HCRT-1 and HCRT-2 (orexin A y B, respectively), synthesized by neurons located in the postero-lateral hypothalamus, whose projections and receptors are widely distributed throughout the brain, including the medial preoptic area (mPOA). HCRT have been associated with a wide range of physiological functions including sleep-wake cycle, maternal behavior and body temperature, all regulated by the mPOA. Previously we showed that HCRT in the mPOA facilitates certain active maternal behaviors, while the blockade of HCRT-R1 increased the time spent in nursing. As mother rats mainly sleep while they nurse, we hypothesize that HCRT in the mPOA of lactating rats reduce sleep and nursing, while the intra-mPOA administration of the dual orexin receptor antagonist (DORA) would generate the opposite effect. Therefore, the aim of this study was to determine the role of HCRT within the mPOA, in the regulation and integration of the sleep-wake cycle, maternal behavior and body temperature of lactating rats. To evaluate this idea, we assessed the sleep-wake states, maternal behavior and body temperature of lactating rats following microinjections of HCRT-1 (100 and 200 μM) and DORA (5mM) into the mPOA. As expected, our data shows that HCRT-1 in mPOA promoted wakefulness and a slightly increase in body temperature, whereas DORA increased both NREM and REM sleep along with nursing and milk ejection. Taken together, our results strongly suggest that the reduction of the endogenous HCRT within the mPOA of lactating rats is important to promote sleep, nursing and milk ejection.

## Introduction

The hypocretinergic (HCRTergic) system has been associated with several physiological functions including the maintenance of wakefulness, the promotion of several motivated behaviors, such as maternal behavior, and the control of body temperature (Boutrel et al., 2010; D’Anna and Gammie, 2006; Harris et al., 2005; McGregor et al., 2011; Muschamp et al., 2007; Rivas et al., 2016; Torterolo et al., 2011; Yoshimichi et al., 2001). In this sense, the sleep disorder called narcolepsy, caused by the degeneration of the hypocretinergic neurons, is characterized by sleep attacks and cataplexy as well as the disruption of thermoregulation (Harding et al., 2019; van der Heide et al., 2016).

The hypocretins (HCRT), also known as orexins, consist of two neuroexcitatory peptides, HCRT-1 and HCRT-2 (also called orexin A y B, respectively), synthesized by neurons located in the postero-lateral hypothalamus, whose projections and receptors are widely distributed throughout the brain, including the medial preoptic area (mPOA) (Peyron et al., 1998; Sakurai et al., 1998; Taheri and Bloom, 2001; Trivedi et al., 1998). HCRT binds to two metabotropic receptors: HCRT type 1 (HCRT-R1) and type 2 receptors (HCRT-R2) with different affinity. HCRT receptors, HCRT-R1 in particular, are expressed in the mPOA (Marcus et al., 2001; Trivedi et al., 1998), a critical region for the regulation of several physiological functions, including sleep and wakefulness, body temperature (Kumar, 2004), as well as the maternal care of the pups (Numan, 1974; Numan and Stolzenberg, 2009; Stolzenberg and Numan, 2011).

There is a great experimental evidence, including studies using different methodological approaches, showing that the mPOA regulates the wake-sleep cycle. Specifically, lesions of the mPOA produced a reduction in non-REM (NREM) and REM sleep (John and Kumar, 1998; Lu et al., 2000; Srividya et al., 2006). In addition, administration of glutamate (Kaushik et al., 2011) or adenosine (Mendelson, 2000) into the mPOA, as well as the activation of GABAergic and galaninergic neurons of this area promote NREM sleep (Chung et al., 2017; Harding et al., 2018; Kroeger et al., 2018; Vanini et al., 2020). Recently, it has been reported that the chemogenetic activation of a group of glutamatergic neurons within the mPOA and surroundings areas increases wakefulness, and decreases both NREM and REM sleep (Mondino et al., 2021; Vanini et al., 2020), suggesting that mPOA contains not only sleep-inducing neurons, but also wake-promoting neurons.

Administration of HCRT-1 into the mPOA of male rats promotes wakefulness and reduces both NREM and REM sleep (Espana et al., 2001), but the effect in lactating rats is unknown. In this sense, there is a great body of evidence showing that the mPOA goes through several changes along the postpartum period (Fleming and Korsmit, 1996; Numan, 2006; Pereira and Morrell, 2009; Rondini et al., 2010; Uriarte et al., 2020). These changes may not only prepare the lactating female to the maternal care of the pups, but also alter the functionality of different circuits that allow to orchestrate maternal behavior with its own physiology. In recent reports, we explored how different neurotransmitters modifies sleep and maternal behavior acting through the mPOA of the lactating rat. Microinjection of the dopamine D2-receptor antagonist Raclopride into the mPOA reduces REM sleep, and its transitional stage from NREM sleep, while NREM sleep is not affected (Benedetto et al., 2017a). On the contrary, the GABA_A_ antagonist Bicuculine has no effect on sleep while increases active maternal care of the pups (Benedetto et al., 2021).

The mPOA also plays an important role in thermoregulation (Srividya et al., 2006). In fact, not only the neural mechanisms that regulates both sleep and body temperature seem to coexist anatomically within the mPOA but are also functionally linked (Cerri and Amici, 2021; Harding et al., 2018; Kumar, 2004). Regarding HCRT, it has been shown that the body temperature is increased by the administration of HCRT-1 into the third ventricle (Yoshimichi et al., 2001) and reduced by HCRT blockade (Martin et al., 2019; Rusyniak et al., 2011). However, its effect within the mPOA is unknown.

The mPOA has been largely known as a key neural site where the hormones and several neuromodulators act to mediate the maternal care of the pups (Benedetto et al., 2014; Rivas et al., 2016; Stolzenberg et al., 2019). Specifically, we have previously showed that HCRT in the mPOA facilitates certain active maternal behavior, while the blockade of HCRT-1 decreases active components of maternal behavior and promotes nursing (Rivas et al., 2016). Since mother rats mainly sleep while they are nursing (Benedetto et al., 2017b), the increased time spent in nursing found with HCRT blockade within the mPOA (Rivas et al., 2016), could be related to a sleep-promoting effect. Therefore, we hypothesize that HCRT administration into the mPOA of lactating rats reduces sleep and nursing, while the blocking of endogenous HCRT provokes the opposite effect. To assess this idea, this study aims to determine the effect of the microinjections of HCRT-1 and the dual orexin receptor antagonist (DORA) into the mPOA of lactating rats on sleep and wakefulness, maternal behavior and body temperature.

## Material and methods

### Animals and housing

Twenty-three primiparous Wistar female rats (250 g) and pups were used in this study. The experimental procedures were in strict accordance with the “Guide for the care and use of laboratory animals” (8th edition. National Academy Press, Washington D. C., 2011) and approved by the Institutional Animal Care Committee (expedient N° 070153-000304). All efforts were made in order to minimize the number of animals and their suffering. We used the same conditions and experimental protocol described in previous articles (Benedetto et al., 2014; Benedetto et al., 2017a; Benedetto et al., 2017b). Animals were housed in a temperature-controlled (22 ± 1 °C) room, under a 12-h light/dark cycle (lights on at 6:00 a.m.), with *ad libitum* access to food and water. Two days before giving birth, pregnant females were housed individually. On postpartum day 1 (PPD1, birth = day 0), litters were culled to four female and four male pups per mother.

### Stereotaxic surgery

Briefly, on PPD1 females were anesthetized with a mixture of ketamine/xylazine/acepromazine maleate (80/2.8/2.0 mg/kg. i.p.). Female rats were bilaterally implanted with 22-gauge stainless steel guide cannulae (Plastic One, Roanoke, VA) aimed 2 mm dorsal to the mPOA: AP −0.5 mm (from Bregma); ML ± 0.5 mm (from midline); DV −6.5 mm (from skull) according to (Paxinos and Watson, 2005). In addition, cortical electroencephalogram (EEG) and dorsal neck muscle electromyogram (EMG) electrodes were implanted for the assessment of sleep and wakefulness (W) states. EEG electrodes were placed in the prefrontal cortex (AP = + 3.0; ML = 2.0), parietal cortex (AP = −4.0, ML = 3.0), occipital cortex (AP = −7.0, ML = 3.0), and over the cerebellum as a reference electrode (AP = −11.0, ML = 0.0). Two additional stainless-steel screws were implanted into the skull as anchors. All electrodes were soldered to a six-pin connector. The connector and the guide cannulae were cemented to the skull using dental acrylic. Immediately after surgery, each mother was reunited with her pups in the home cage.

At the end of the stereotaxic surgery, the lower dorsal part of the flank was shaved and a 1.5–2 cm long incision was made on the skin. A temperature data logger iButton (Thermochron, model DS2422) inside a silicone-coat capsule was implanted subcutaneously; thereafter, the skin was sutured. The iButton was implanted in the dorsal part of the animal to avoid interference with the mammary glands and suckling of the pups.

A pre-surgery single dose of ketoprofen (5mg/kg, s.c.) was used to reduce pain. Also, topic antibiotic (Crema 6A, Labyes) was applied into the brain surgery injury and abdominal incision, and sterile 0.9 % saline (10 ml/kg, s.c.) was administrated post-surgery to prevent dehydration during recovery (Benedetto et al., 2021).

### Experimental design

All experiments were performed between PPD4-8 during the light phase. Before experiments began, a baseline recording session was performed to corroborate that all sleep parameters and maternal behaviors were adequate. Animals were randomly assigned to one of the following independent groups: HCRT or DORA group. For the HCRT group, each animal received a total of three microinjections: HCRT-1 100 μM (HCRT_100_), HCRT-1 200 μM (HCRT_200_) and sterile saline as vehicle. For DORA group, each animal received two microinjections: DORA 5 mM and dimethyl sulfoxide (DMSO) as vehicle. All microinjections were administrated in a counterbalanced design and the day after the microinjections, no experiments were performed.

### Drugs

HCRT-1 (BACHEM, Bubendorf, Switzerland) was diluted in sterile saline to obtain a final concentration of 100 and 200 μM. Aliquots for these doses were prepared in advance, frozen at −20°C, and thawed immediately before use. DORA TCS-1102 (Sigma–Aldrich, St Louis, USA) was diluted in DMSO 10% to obtain a final concentration of 5 mM. Similar doses were used in previous studies (Hsiao et al., 2012; Korim et al., 2014; Rivas et al., 2016).

### Microinjection procedure

Microinjection procedure was performed using the injection cannulae (28 gauge; Plastic One, Roanoke, VA) extending 2 mm beyond the tip of the guide cannulae, and a constant-rate infusion pump (Harvard apparatus, USA). The rats were bilaterally microinjected with 0.2 μl of either HCRT-1, DORA, or the same volume of the correspondent vehicle into the mPOA over a period of 2 min. The injection cannulae were left in place for an additional minute to allow for the diffusion of the drug. The same microinjection volume was used in previous studies of the group (Benedetto et al., 2014; Benedetto et al., 2017a; Benedetto et al., 2021; Rivas et al., 2016).

### Experimental sessions

During each experimental day, at 9 a.m., pups were removed from the maternal cage for three hours and placed under a heat lamp. Fifteen minutes before the completion of maternal separation, microinjection procedure was performed. Thereafter, the rat was returned to her home cage and connected to the recording system. Then, the entire litter was weighed, and at the completion of the separation period the pups were scattered in the mothers’ home cage opposite to the nest. Subsequently, polysomnographic recording and videotaping of the maternal behavior was initiated for four hours. After each recording session, the mother rat was disconnected from the recording device and the entire litter was weighed again.

### Sleep recording

Bioelectric signals were amplified (×1000), filtered (0.1–500 Hz), sampled (1024 Hz, 16 bits) and stored in a PC for further analysis using the Spike 2 software. The states of light sleep (LS), slow wave sleep (SWS), REM sleep and W were determined in 5-s epochs with standard criteria (Benedetto et al., 2017a; Benedetto et al., 2017b; Benedetto et al., 2013). Additionally, the intermediate stage (IS, transition from NREM to REM sleep) was also distinguished (sleep spindles combined with theta activity) (Gottesmann, 1992). Total time spent in W, LS, SWS, NREM sleep (LS + SWS), IS and REM sleep over the total recording time and each hour separately were analyzed. In addition, sleep latencies (first episodes ≥ 20 s from the beginning of the recordings), number, and duration of episodes of each state were studied.

### EEG spectral power analysis

EEG analysis of power (1 – 45 Hz) was conducted during the 4 hours after delivery of vehicle or drugs into mPOA. Spectrograms (time-frequency representation of the EEG signal) were examined in Spike 2. To further study the spectral characteristics, raw EEG signals from prefrontal and parietal channels were exported into MATLAB. The average power spectrum was obtained by means of the *pwelch* built-in MATLAB function (parameters: window = 1024, noverlap = [], fs = 1024, nfft = 1024), which corresponds to 1-s sliding windows with half-window overlap, and a frequency resolution of 1 Hz. All spectra were normalized to obtain the relative power by dividing the power value of each frequency by the sum across frequencies. Because notch filters (50 Hz) were applied to EEG channels in some recordings, we only considered the frequency bands up to 45 Hz. For each animal in each treatment group, the mean power spectrum in each behavioral state (W, LS, SWS, IS, REM) was obtained by averaging the power spectra across all available windows in prefrontal and parietal cortices.

### Maternal behavior

Maternal behavior was analyzed from digital videos and classified into three major categories: hovering over the pups (dam over the pups while actively engaged in any activity), nursing (low and high kyphosis and supine postures), and away from the pups. Maternal states were staged in 5-s epochs and analyzed for the 4-h-recording session taking each hour separately.

In addition, specific active maternal behaviors were measured: the number of retrievals of the pups into the nest and the latency to group the entire litter. Besides, the latency to the first nursing bout ≥ 2 min, the number of milk ejections (indirectly through the stretching behavior of the pups (Lincoln et al., 1973; Voloschin and Tramezzani, 1979) and the percentage of the litter weight gain (as an indirect measurement of the amount of ejected milk (Benedetto et al., 2021; Lincoln et al., 1973; Peña et al., 2020; Stern, 1991) were recorded.

### Temperature recording

The body temperature was automatically recorded by the iButtons. The onset of measurement was set to initiate 3 days post-surgery for a 5-day period, taking temperature readings every 3 minutes. The temperature resolution of the iButtons was 0.0625°C. After the experiments were completed and the animals were sacrificed, the iButtons were removed and data acquired were downloaded to a PC.

### Histological verification of microinjection sites

At the end of the experiment the animals were euthanized with an overdose of ketamine/xylazine, perfused with 4% paraformaldehyde, and their brains were removed for histological processing. Thereafter, the brains were cut in 100 μm coronal sections with a vibratome. The location of mPOA microinjection sites were verified according to the neuroanatomical atlas of Paxinos and Watson (Paxinos and Watson, 2005).

### Statistics

All values are presented as mean ± S.E.M (standard error). Comparisons of sleep and maternal parameters among HCRT-1 groups were evaluated by one-way repeated measures (ANOVA) followed by Tukey post hoc test, while paired Student t-test was used to compare DORA groups. Differences in EEG power among groups were evaluated utilizing the Friedman test for the HCRT-1 group, and Wilcoxon signed-rank test for the DORA group, since the data did not show a normal distribution tested with the Lilliefors test. The criterion used to discard the null hypotheses was p < 0.05.

## Results

### Sites of injection

As depicted in Figure 1, all microinjections included in the study were located within the mPOA between −0.12 and −0.60 mm from Bregma, based on the examination of the cannulae tracks in the histological sections (Paxinos and Watson, 2005). In six animals the guide cannulae was located outside the mPOA and were excluded from the data analysis. Therefore, a total of 16 animals were included in the study. Figure 1 shows the microinjection sites of HCRT-1 and DORA groups.

**Figure 1.**
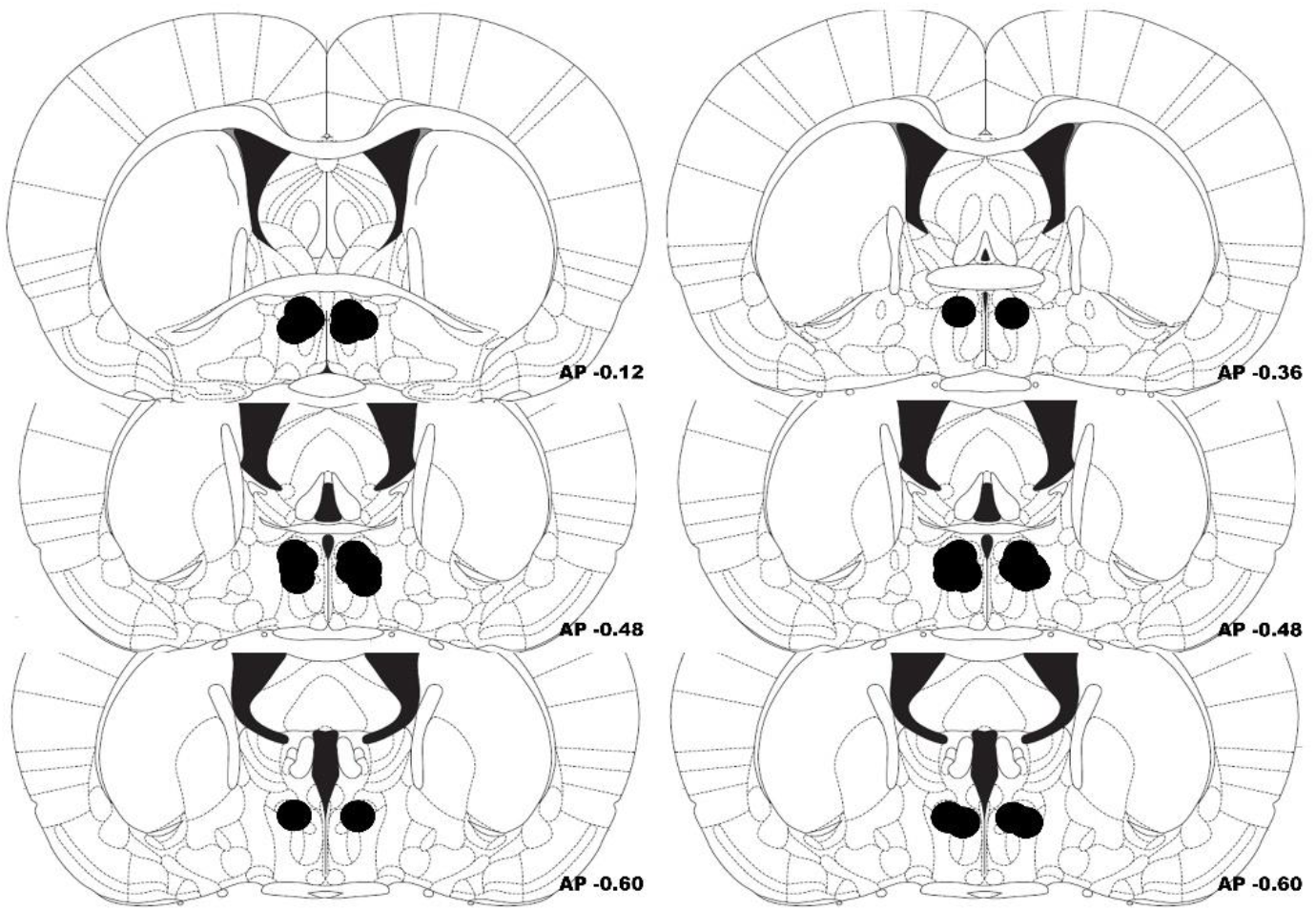
Microinjections sites. Schematic representations of coronal sections at the level of mPOA. Black circles indicate the microinjection sites of the HCRT-1 (left) and DORA group (right); bottom numbers indicate distance from Bregma. Plates were taken from the atlas of (Paxinos and Watson, 2005).

### Effect of HCRT-1 on sleep and waking states

Representative hypnograms and spectrograms (time-frequency representation of the EEG signal) from 0 to 30 Hz is shown for the HCRT group in Figure 2A. Sleep and waking parameters means are shown in Table 1 and Figures 2A and 3. Compared to vehicle, the total time spent in W significantly increased after the delivery of both HCRT_100_ and HCRT_200_. Furthermore, HCRT_200_ significantly decreased the total time spent in SWS, and there was a tendency to reduce the duration of the SWS episodes (Table 1 and Figures 2A and 3).

**Table 1.** Effects of microinjection of HCRT-1 into mPOA on sleep parameters during 4-hour sessions.

|  | Vehicle | HCRT<sub>100</sub> | HCRT<sub>200</sub> | ANOVA |  | Tukey |
| --- | --- | --- | --- | --- | --- | --- |
|  |  |  |  | F | P | P |
| **Wakefulness** |  |  |  |  |  |  |
| Total duration (min) | 105.29 ± 8.01 | 117.43 ± 6.06* | 120.74 ± 6.15* | 7.64 | 0.005 | 0.001<br>0.003 |
| Number of episodes | 133.44 ± 8.32 | 145.33 ± 8.74 | 155.56 ± 12.40 | 1.51 | 0.250 |  |
| Episodes duration (min) | 0.81 ± 0.09 | 0.83 ± 0.08 | 0.80 ± 0.08 | 0.09 | 0.911 |  |
| **Light Sleep** |  |  |  |  |  |  |
| Total duration (min) | 32.19 ± 2.62 | 32.00 ± 2.55 | 34.04 ± 2.94 | 0.31 | 0.737 |  |
| Number of episodes | 186.44 ± 12.64 | 197.44 ± 14.87 | 209.56 ± 19.34 | 1.28 | 0.305 |  |
| Episodes duration (min) | 0.17 ± 0.01 | 0.16 ± 0.01 | 0.16 ± 0.01 | 0.93 | 0.414 |  |
| **Slow Wave Sleep** |  |  |  |  |  |  |
| Total duration (min) | 83.17 ± 4.78 | 73.20 ± 5.28 | 73.03 ± 4.69* | 4.41 | 0.030 | 0.049 |
| Number of episodes | 133.78 ± 9.61 | 139.11 ± 11.22 | 143.78 ± 13.33 | 0.63 | 0.544 |  |
| Episodes duration (min) | 0.63 ± 0.03 | 0.54 ± 0.06 | 0.52 ± 0.05 | 3.91 | 0.041 | 0.051 |
| **Intermediate stage** |  |  |  |  |  |  |
| Total duration (min) | 5.22 ± 1.34 | 6.39 ± 1.15 | 4.66 ± 1.29 | 0.85 | 0.445 |  |
| Number of episodes | 16.33 ± 2.99 | 19.33 ± 3.02 | 12.67 ± 3.40 | 2.87 | 0.086 |  |
| Episodes duration (min) | 0.30 ± 0.05 | 0.32 ± 0.05 | 0.31 ± 0.06 | 0.09 | 0.915 |  |
| **REM Sleep** |  |  |  |  |  |  |
| Total duration (min) | 14.13 ± 2.09 | 10.98 ± 1.71 | 7.54 ± 1.54* | 10.04 | 0.001 | 0.001 |
| Number of episodes | 12.44 ± 2.00 | 9.89 ± 1.48 | 6.33 ± 1.73* | 8.67 | 0.003 | 0.002 |
| Episodes duration (min) | 1.23 ± 0.23 | 1.18 ± 0.20 | 1.21 ± 0.31 | 0.04 | 0.964 |  |
| **Latency NREM (min)** | 13.13 ± 2.29 | 13.25 ± 4.10 | 18.90 ± 3.82 | 1.81 | 0.195 |  |
| **Latency REM (min)** | 81.41 ± 11.88 | 91.90 ± 11.05 | 136.94 ± 19.90* | 7.26 | 0.006 | 0.007 |

Data is presented as mean ± standard error of nine rats. \* denote significant difference compared to control values, using one-way repeated measures ANOVA followed by Tukey test.

**Figure 2.**
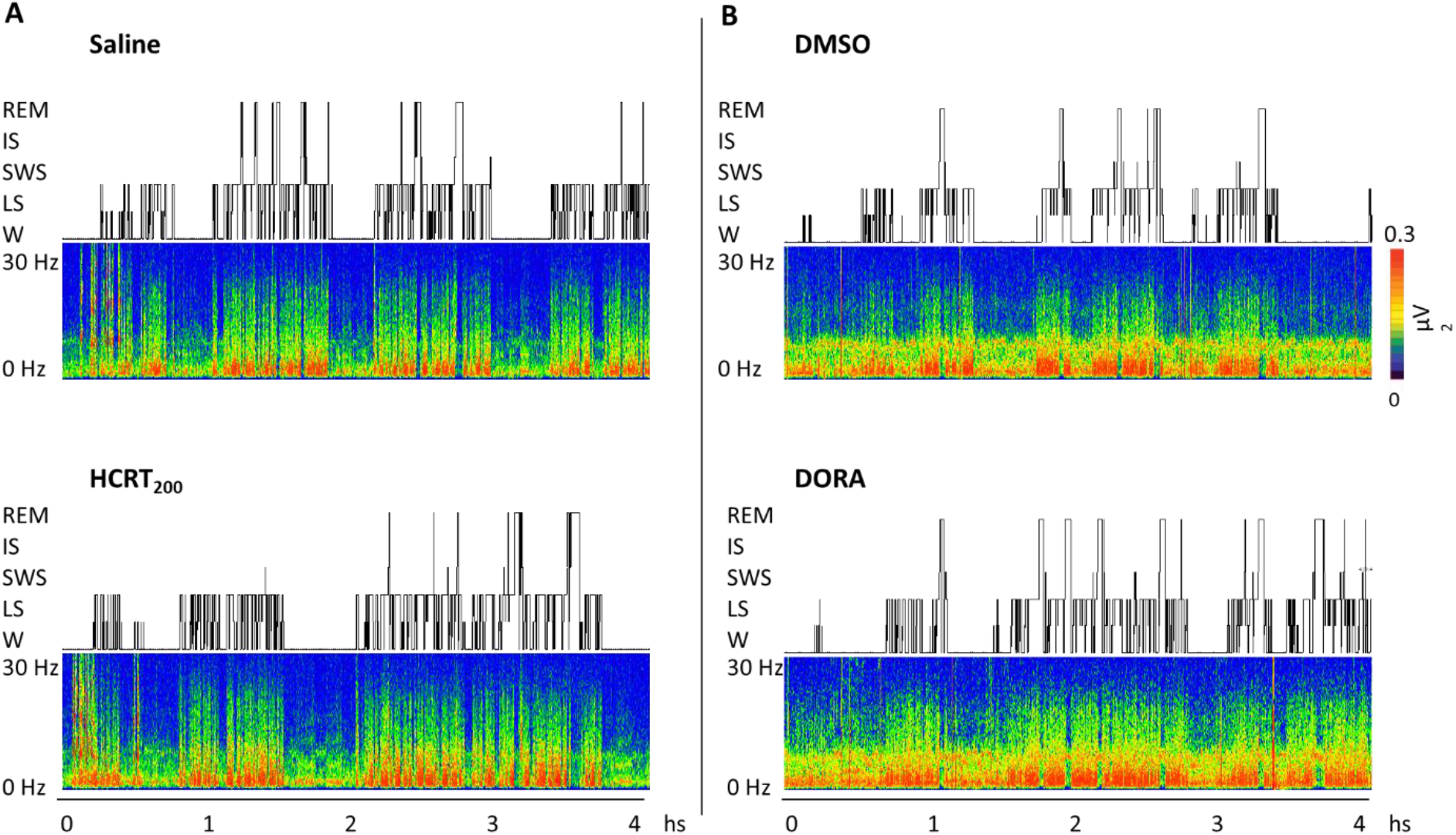
Hypnograms and spectrograms (1 - 30 Hz) from the prefrontal cortical recordings of a representative animal are shown after saline and HCRT_200_ (A), and DMSO and DORA (B) local administration. W, wakefulness; LS, light sleep; SWS, slow wave sleep; IS, intermediate stage and REM, rapid eyes movements sleep.

**Figure 3.**
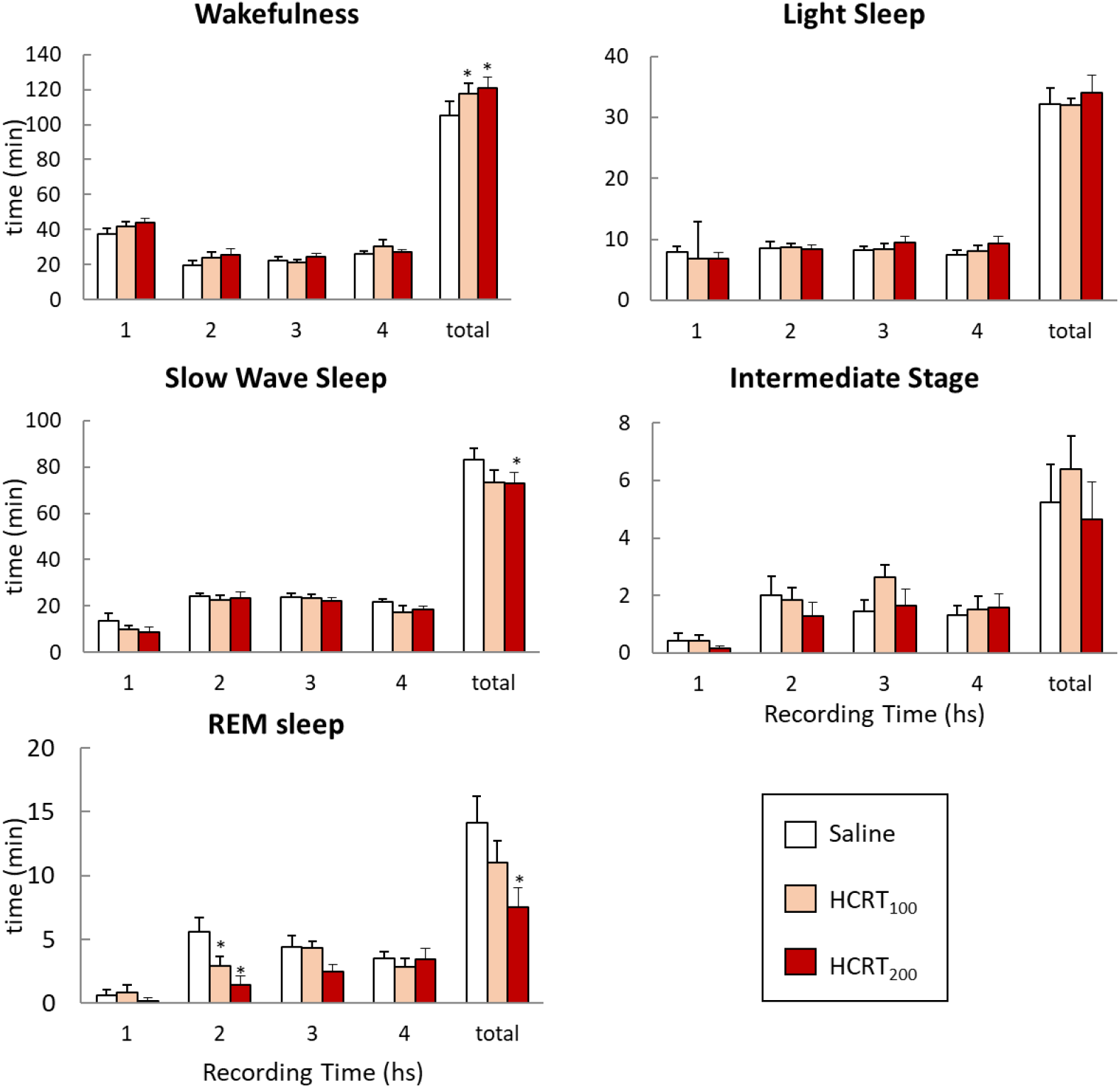
Effect of HCRT-1 microinjection into mPOA on sleep and wakefulness. The charts show the mean time (± SEM) spent in wakefulness, light sleep, slow wave sleep, intermediate stage and REM sleep after local administration of saline, HCRT_100_ and HCRT_200_ during each hour and the total recording time. Group differences were determined by one-way repeated measures ANOVA followed by Tukey as post hoc; asterisks (*) indicates significant differences compared to control values.

Different REM sleep parameters were modified after HCRT. Specifically, the total time spent in REM was significantly reduced following HCRT_200_ microinjections. Also, the time spent in REM during the second hour was different among groups (F (2, 16) = 11.11, p = 0.001). Specifically, it was significantly reduced after local delivery of HCRT_100_ (p = 0.023) and HCRT_200_ (p = 0.001) compared to vehicle (see Figures 2A and 3). Also, the number of REM episodes decreased and latency to REM increased after HCRT_200_ when compared to control values (Table 1).

As shown in Table 1 and Figure 3, no other sleep parameter differed among groups.

### Effect of DORA on sleep and waking states

Representative hypnograms and spectrograms is shown for the DORA group in Figure 2B. The total time spent in W, as well as the time spent in this state during the fourth recording hour (*t* = 2.75, p = 0.016), were significantly reduced after microinjection of DORA into mPOA compared to vehicle (Table 2 and Figure 4). In addition, there was a significant reduction in the duration of W episodes (Table 2). In concordance, DORA local delivery increased the total time spent in SWS as well as the time spent during the fourth recording hour (*t* = 2.62, p = 0.034; Figures 2B and 4).

**Table 2.** Effects of microinjection of DORA into mPOA on sleep parameters during 4-hour sessions.

|  | DMSO | DORA | t | P |
| --- | --- | --- | --- | --- |
| **Wakefulness** |  |  |  |  |
| Total duration (min) | 131.61 ± 5.63 | 115.88 ± 3.08* | 3.10 | 0.017 |
| Number of episodes | 109.25 ± 12.19 | 127.88 ± 10.30 | 2.16 | 0.068 |
| Episodes duration (min) | 1.32 ± 0.18 | 0.95 ± 0.08* | 2.61 | 0.035 |
| **Light Sleep** |  |  |  |  |
| Total duration (min) | 27.75 ± 3.80 | 28.54 ± 3.06 | 0.34 | 0.743 |
| Number of episodes | 159.75 ± 16.23 | 171.63 ± 13.12 | 0.10 | 0.351 |
| Episodes duration (min) | 0.17 ± 0.01 | 0.16 ± 0.01 | 0.86 | 0.416 |
| **Slow Wave Sleep** |  |  |  |  |
| Total duration (min) | 67.67 ± 4.64 | 78.26 ± 3.16* | 3.68 | 0.008 |
| Number of episodes | 110.25 ± 7.01 | 120.00 ± 5.18 | 1.22 | 0.262 |
| Episodes duration (min) | 0.62 ± 0.04 | 0.66 ± 0.03 | 0.78 | 0.461 |
| **Intermediate stage** |  |  |  |  |
| Total duration (min) | 4.83 ± 1.53 | 5.38 ± 1.08 | 0.63 | 0.550 |
| Number of episodes | 15.00 ± 3.24 | 15.62 ± 4.11 | 0.22 | 0.822 |
| Episodes duration (min) | 0.31 ± 0.03 | 0.38 ± 0.04 | 1.69 | 0.135 |
| **REM Sleep** |  |  |  |  |
| Total duration (min) | 8.14 ± 1.22 | 11.94 ± 1.19* | 2.57 | 0.037 |
| Number of episodes | 7.75 ± 1.61 | 8.87 ± 2.18 | 0.59 | 0.575 |
| Episodes duration (min) | 1.35 ± 0.30 | 1.71 ± 0.25 | 1.33 | 0.225 |
| **Latency REM (min)** | 86.43 ± 15.53 | 65.33 ± 7.38 | 1.95 | 0.091 |
| **Latency NREM (min)** | 14.40 ± 3.86 | 9.73 ± 0.99 | 1.18 | 0.276 |

Data is presented as mean ± standard error of eight rats. \* denote significant difference compared to control values, using Student paired test.

**Figure 4.**
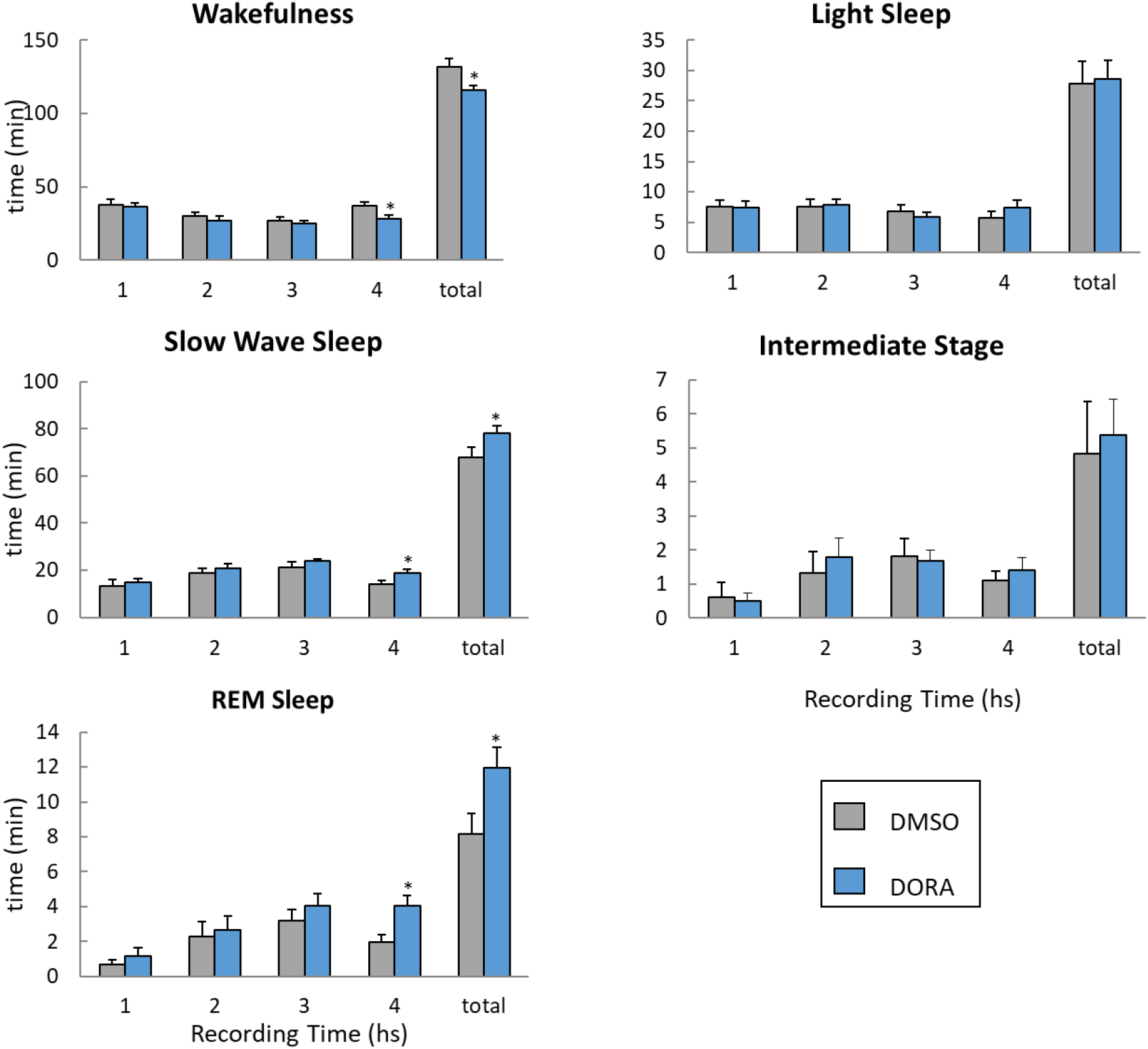
Effect of DORA microinjection into mPOA on sleep and wakefulness. The charts show the mean time spent in wakefulness, light sleep, slow wave sleep, intermediate stage and REM sleep after local administration of vehicle (DSMO 10%) and DORA 5 mM during each hour and the total recording time. Group differences were determined by paired Student test; * indicates significant differences compared to control values.

Besides, the total time spent in REM increased after DORA microinjection, as well as the time in REM during the fourth recording hour (*t* = 3.12, p = 0.007; Figures 2B and 4). Sleep parameters of LS and IS were not affected by DORA microinjection (Table 2 and Figure 4).

### EEG spectral analysis following administration of HCRT-1 and DORA

An example spectrogram (time-frequency representation of the EEG signal) from 0 to 30 Hz for each experimental group is shown in Figure 4 and the mean power in Figure 5. Compared to vehicle, following HCRT-1 or DORA microinjections into the mPOA the EEG power did not differ in any frequency studied, neither in prefrontal, parietal cortex, W or sleep (Figure 5). We also extended the frequency bands studied up to 200 Hz, but no differences were found (data not shown).

**Figure 5.**
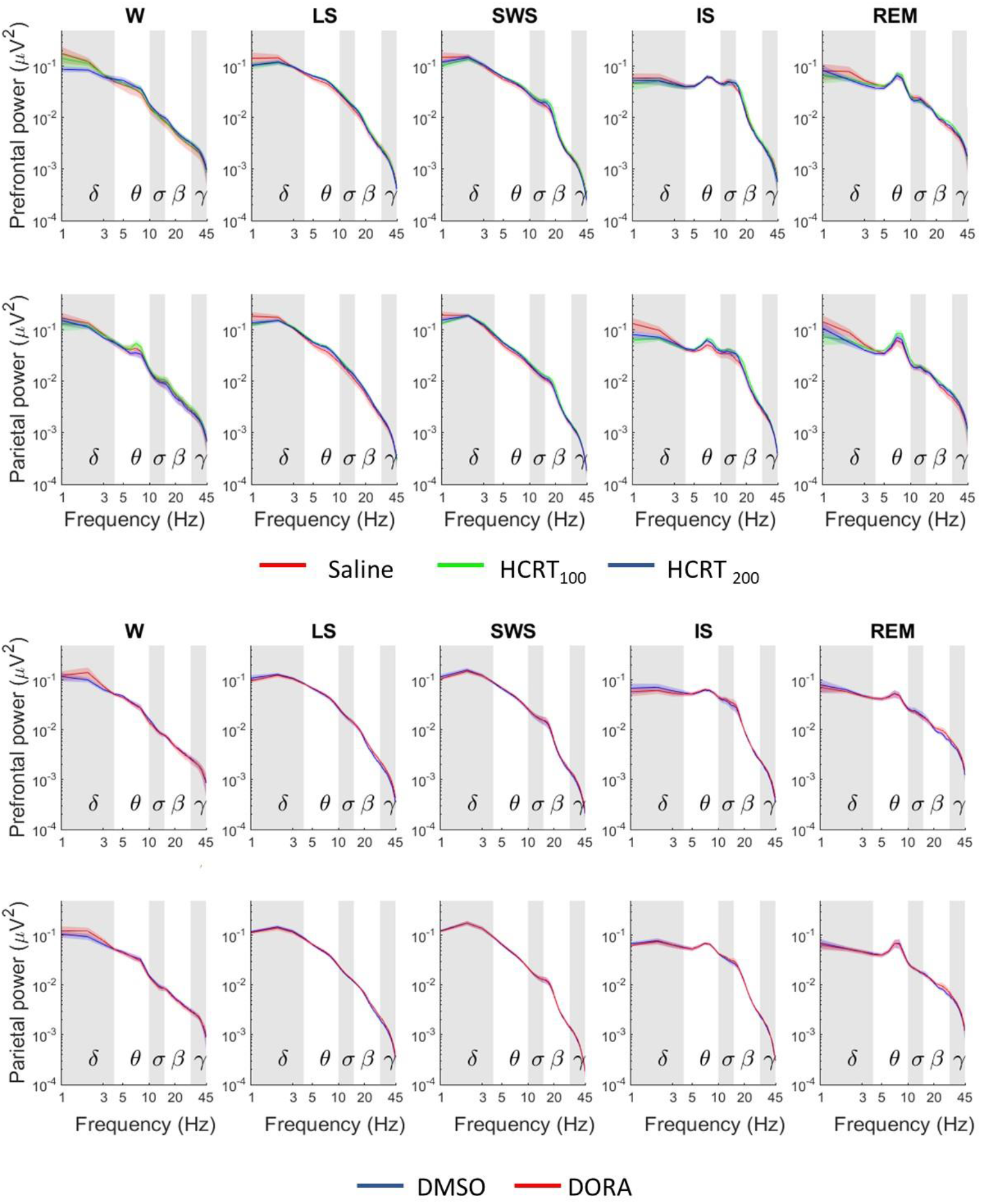
The graphs plot spectral power changes in the prefrontal and parietal cortex during every sleep state for frequencies between 1 and 30 Hz for HCRT-1 (top) and DORA (bottom) groups. Traces represent mean values (thin, dark lines) ± SEM (shaded area above and below the mean). Frequency ranges are indicated by alternating horizontal colored bands in the background of the graphs: Delta (1-4 Hz), Theta (4-10 Hz), Sigma (10-15 Hz), Beta (15-30 Hz) and Gamma (30-45 Hz). For HCRT-1 Friedman test for multiple comparisons was employed for statistical comparison of spectral power in each frequency with control. Wilcoxon signed-rank test was used for DORA group. W, wakefulness; LS, light sleep; SWS, slow wave sleep; IS, intermediate stage and REM, rapid eyes movements sleep.

### Effect of HCRT-1 on maternal behavior

Table 3 shows the results of HCRT-1 microinjection into mPOA on maternal behavior parameters. Only the litter weight gain decreased following HCRT_100_ microinjection. HCRT-1 did not produce any additional significant changes in the maternal behaviors analyzed (Figure 6).

**Table 3.** Effects of microinjection of HCRT-1 into mPOA on maternal behavior parameters during 4-hour sessions.

|  | Vehicle | HCRT<sub>100</sub> | HCRT<sub>200</sub> | ANOVA |  | Tukey |
| --- | --- | --- | --- | --- | --- | --- |
|  |  |  |  | F | P | P |
| *Latency to (min):* |  |  |  |  |  |  |
| Reunion litter | 8.50 ± 4.11 | 14.20 ± 5.00 | 8.75 ± 4.05 | 0.62 | 0.549 |  |
| Nursing | 8.69 ± 1.29 | 12.10 ± 3.92 | 8.74 ± 1.40 | 0.54 | 0.593 |  |
| *Duration (min):* |  |  |  |  |  |  |
| Nursing total | 171.52 ± 6.99 | 162.46 ± 7.61 | 154.43 ± 12.37 | 1.70 | 0.214 |  |
| Nursing Episodes | 12.44 ± 1.25 | 10.50 ± 0.78 | 11.29 ± 1.61 | 0.78 | 0.476 |  |
| *Number of:* |  |  |  |  |  |  |
| Nursing episodes | 15.11 ± 1.31 | 15.78 ± 0.70 | 15.56 ± 1.58 | 0.07 | 0.928 |  |
| Milk ejections | 21 ± 2.09 | 19.67 ± 2.06 | 16.67 ± 1.29 | 2.41 | 0.121 |  |
| Litter weight gain (%) | 7.08 ± 0.72 | 4.77 ± 0.49* | 5.21 ± 0.57 | 4.16 | 0.035 | 0.038 |

Data is presented as mean ± SEM of nine rats. \* denote significant difference compared to control values, using one-way repeated measures ANOVA followed by Tukey test.

**Figure 6.**
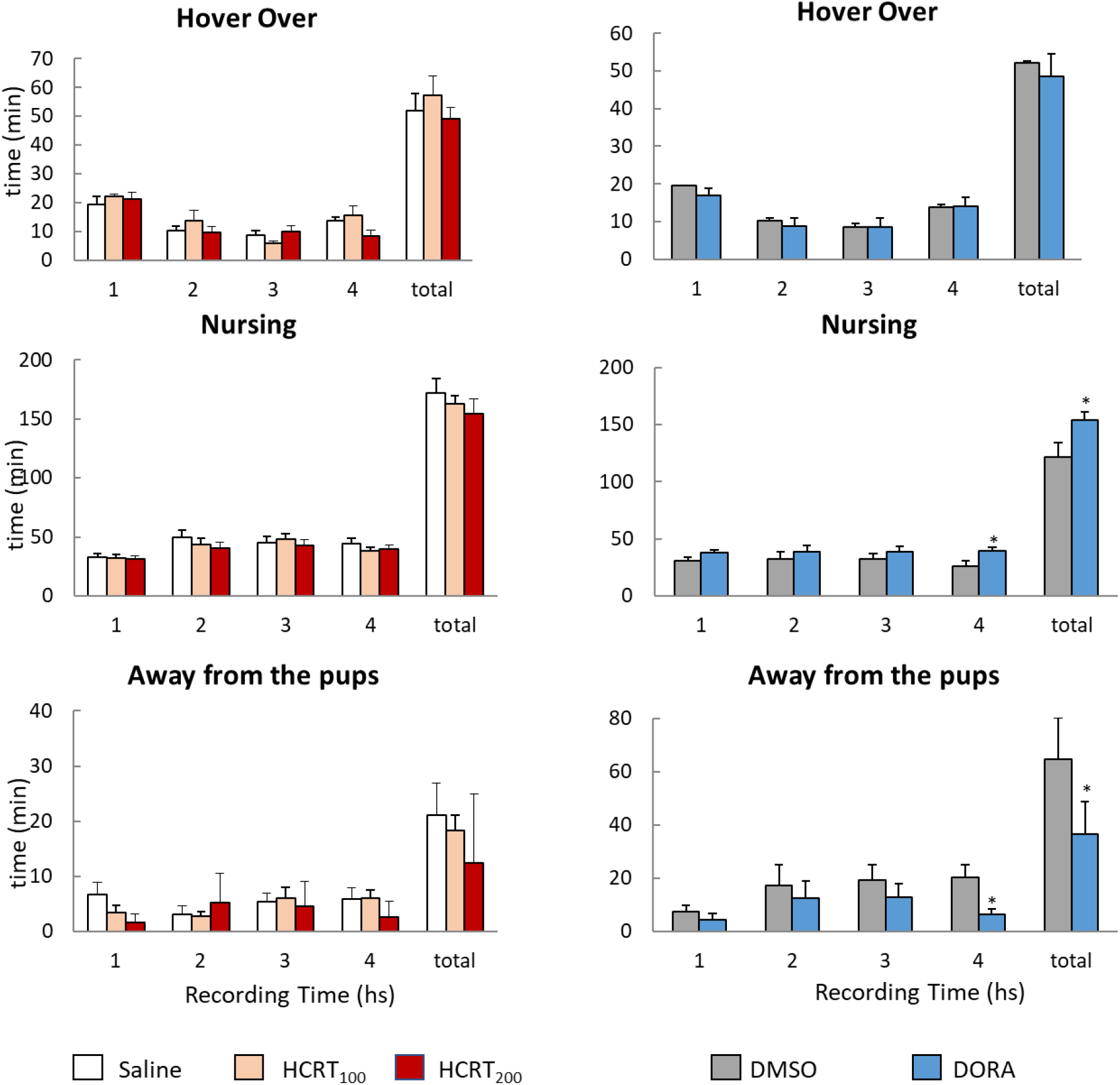
Effect of HCRT-1 and DORA microinjection into mPOA on maternal behaviors. The charts show the mean time spent in hover over, nursing, and away from the pups after local administration of saline, HCRT_100_ and HCRT_200_ (on the left), and DMSO and DORA (on the right), during each hour and the total recording time. Group differences were determined by one-way repeated measures ANOVA followed by Tukey in HCRT-1 groups, and Student paired test in DORA group; * indicates significant differences compared to control values.

### Effect of DORA on maternal behavior

Compared with vehicle administration, the local delivery of DORA into the mPOA produced an increase in the time that females spent nursing their pups. This enhancement was observed in the total recording time (Table 4) as well as during the fourth recording hour (*t* = 2.40, p = 0.034; Figure 6). In accordance, the number of milk ejections increased after microinjection of DORA compared to vehicle, and litter weight gain tended to increase (Table 4). Furthermore, DORA local delivery decreased the total time that dams spent away from the pups (*t* = 4.27, p = 0.005; Figure 6), as well as during the fourth recording hour (*t* = 4.81, p = 0.003; Figure 6). There were no additional significant changes in other maternal behavior parameters analyzed (Table 4).

**Table 4.** Effects of microinjection of DORA into mPOA on maternal behavior parameters during 4-hour sessions.

|  | DMSO | DORA | t-test | P |
| --- | --- | --- | --- | --- |
| *Latency to (min):* |  |  |  |  |
| Reunion litter | 1.56 ± 0.09 | 9.75 ± 5.24 | 1.55 | 0.172 |
| Nursing | 7.71 ± 1.46 | 6.12 ± 0.72 | 0.87 | 0.418 |
| *Duration (min):* |  |  |  |  |
| Nursing total | 121.38 ± 12.67 | 154.15 ± 6.85* | 4.07 | 0.007 |
| Nursing Episodes | 10.01 ± 0.78 | 12.67 ± 2.05 | 0.10 | 0.356 |
| *Number of:* |  |  |  |  |
| Nursing episodes | 12.14 ± 0.86 | 14 ± 2.12 | 1,30 | 0,239 |
| Milk ejections | 17 ± 2.31 | 27.14 ± 3.39* | 4.33 | 0.005 |
| Litter weight gain (%) | 4.50 ± 0.57 | 5.32 ± 0.50 | 2.09 | 0.071 |

Data is presented as mean ± standard error of seven rats, \* denote significant difference between groups, using Student paired test.

### Effect of HCRT-1 and DORA on body temperature

Compared to vehicle microinjections, body temperature differs among groups. Specifically, it increased after microinjection of HCRT_200_ during the first (F (2, 14) = 9.27, p = 0.004; Tukey: HCRT_200_ vs. saline p = 0.027) and second hour F (2, 14) = 6.83, p = 0.010; Tukey: HCRT_200_ vs. saline p = 0.017) compared to control values (Figure 7).

**Figure 7.**
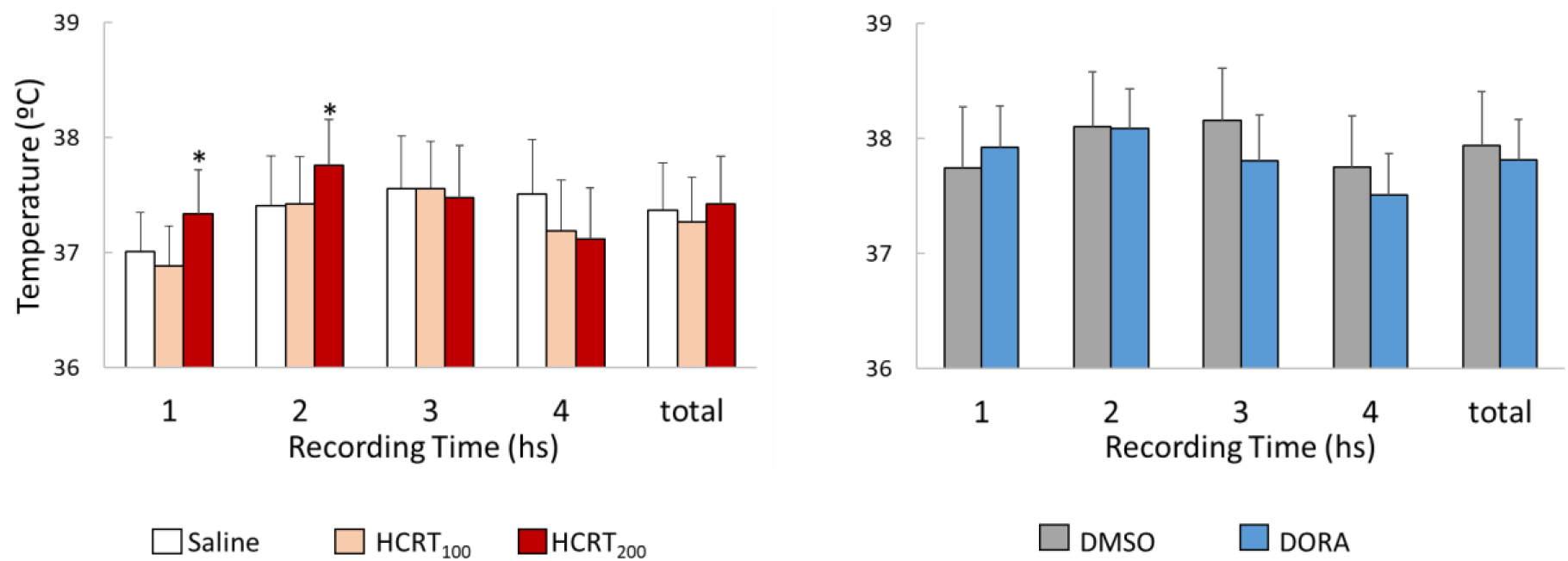
Effect of HCRT-1 and DORA microinjection into mPOA on body temperature. The charts show the average ± SEM body temperature after local administration of saline, HCRT_100_ and HCRT_200_ (on the left), and DMSO and DORA (on the right), during each hour and in the total recording time. Group differences were determined by one-way repeated measures ANOVA followed by Tukey in HCRT-1 groups, and Student paired test in DORA group. * indicates significant differences compared to control values.

DORA microinjection in the mPOA did not provoke significant changes in the body temperature of mother rats, neither in the total recording time nor during each hour analyzed separately (Figure 7).

## Discussion

This study shows that local perfusion of HCRT-1 into mPOA of lactating rats increased total recording time spent awake and decreased time in both SWS and REM sleep, while the dual receptor antagonist of HCRT-receptors or DORA had the opposite effect. Together with the enhancement of sleep, local administration of DORA increased the time spent in nursing the pups and the number of milk ejections. These facts suggest that sleep and nursing, can be promoted together. In addition, while HCRT-1 microinjections increased body temperature, this parameter was not affected by DORA.

### Technical considerations

We performed intracerebral microinjection of HCRT-1 and DORA in a volume of 0.2 μl, similarly to previous studies of our laboratory (Benedetto et al., 2014; Benedetto et al., 2017a; Benedetto et al., 2013; Lagos et al., 2009; Lagos et al., 2011; Rivas et al., 2016). Since the same volume of methylene blue has been shown to diffuse approximately 500 μm in the CNS (Lohman et al., 2005), we cannot rule out the possibility that the drugs could have potentially affected limits of adjacent areas, as the lateral preoptic area, which are also a somnogenic areas.

### HCRT-1 microinjections into the mPOA modifies sleep but not EEG activity

The fact that HCRT-1 infusion into the mPOA of lactating mother rats reduced sleep is consistent with previous evidence in male rats showing that microinjections of HCRT-1 in the preoptic area (POA) promotes W, in spite that the target of the injection was more lateral than in the present study (Espana et al., 2001; Methippara et al., 2000). Taken together, we can hypothesize that HCRT-1 function on sleep is widespread along the POA and not restricted to the medial zone of the mPOA, and is consistent both in male and mother rats.

In accordance with HCRT-1 effects, our results show that local infusion of DORA decreased total time spent awake while increased the time in SWS and REM sleep. These data strongly suggest that there is a tonic endogenous release of HCRT in mPOA sleeping circuits. It is important to note, that the DORA Suvorexant has been approved in 2014 as a hypnotic drug by the Food Drug Administration (FDA) (Yang, 2014). Nevertheless, to the best of our knowledge, its effect during the postpartum period has not been studied before, neither in humans nor in animal models.

The mPOA is a heterogeneous region composed of cells that release several neurotransmitters, such as GABA, glutamate, dopamine, and several neuropeptides (Simerly et al., 1986; Tsuneoka et al., 2013). Although most of the neurons whose activity have been related to sleep are GABAergic (Fang et al., 2018; Lonstein and De Vries, 2000; Tsuneoka et al., 2013), it has been recently showed that a subgroup of glutamatergic neurons within the mPOA promotes NREM sleep together with body cooling (Harding et al., 2018). In addition to somnogenic POA neurons, it has been recently reported a group of glutamatergic neurons within the mPOA whose chemogenetic activation increases wakefulness, and decreases both NREM and REM sleep (Mondino et al., 2021; Vanini et al., 2020). Regarding the mechanisms underlying the effects of HCRT in POA neurons, (Eggermann et al., 2001) have studied the effects of HCRT-1 perfusion in neurons of VLPO in rat brain slices. Specifically, they show that HCRT have no effects on the GABA sleep-promoting neurons of the ventrolateral preoptic area (VLPO), whereas they have a strong and direct excitatory effect on the cholinergic neurons of the adjacent basal forebrain, and this effect was dependent on the activation of the HCRT-R2. Moreover, in median preoptic nucleus (MnPO) neurons, through patch-clamp recording in rat brain slices, (Kolaj et al., 2008) have shown that HCRT applications induced a direct postsynaptic depolarization and excitation of glutamatergic currents but had no influence on GABAergic currents. Although sleep active and sleep promoting neurons of the POA are present mainly in the VLPO and MnPO (Gong et al., 2004; Kroeger et al., 2018; Sherin et al., 1996; Szymusiak et al., 1998; Vanini et al., 2020), somnogenic neurons are also present in mPOA (Chung et al., 2017; Harding et al., 2018). However, there are no previous reports exploring the effect of HCRT in mPOA neurons. Although the present study does not provide information about the specific neurons that were affected by HCRT-1 to produce the observed effects, given previous evidence we can speculate that HCRT-1 local administration into the mPOA could be acting directly into glutamatergic rather than in GABAergic neurons.

Even though HCRT-1 and DORA in mPOA affected sleep-wake times, these changes were not accompanied with EEG power modifications in any frequency or behavioral stage studied. This is in accordance with (Hungs and Mignot, 2001), who showed that HCRT neuron ablation in mice not affected the EEG power spectrum in any state. However, intracerebroventricular HCRT-1 administration in male rats decreased delta and alpha power, and theta and beta power increased (Toth et al., 2012); while a low dose of HCRT-1 (140 pmol) only decreased delta power but had no changes in theta-potency (Magdaleno-Madrigal et al., 2019). Collectively, these data suggest that HCRT-1 effects on EEG power are dependent on other brain areas rather than the mPOA. Interestingly, in humans, EEG power spectra in non-REM sleep was not affected by the hypnotic DORA SB-649868 (Bettica et al., 2012); while Suvorexant at clinically effective doses has limited effects on power spectral density, suggesting that the antagonism of the HCRT pathway might lead to improvements in sleep without major changes in the patient’s EEG profile (Ma et al., 2014).

### HCRT into the mPOA modifies maternal behavior

Our results show that the blockade of the action of endogenous HCRT intra-mPOA by DORA increased the time that mother rats spent nursing and the number of milk ejections. The increase in nursing occurred at the expense of a decrease in the time that dams spent away from the pups without affecting the time in hovering over the pups, indicating that DORA increased the time that mothers spent with the pups. This is in accordance with (Grieb et al., 2018), who showed that the endogenous levels of HCRT-1 within the mPOA are negatively correlated with the frequency of contact with the litter and kyphosis postures. However, hover over the pups was negatively correlated with the high levels of HCRT-1. These results together suggest that natural differences in endogenous HCRT levels within the mPOA may lead to maternal behavior differences among individuals.

In contrast, most maternal behaviors analyzed in the present report were undisturbed by the administration of HCRT-1 into mPOA. Only HCRT_100_ reduced litter weight gain, but without affecting nursing time. This is consistent with our previous study, in which latencies and durations of nursing behaviors did not differ after microinjections of HCRT-1 10 and 100 μM, measured in a 30-minute maternal test (Rivas et al., 2016). Together, this lack of effect of HCRT-1 administration on maternal behaviors suggests that the endogenous HCRT tonic levels in mPOA might be already high. Hence, HCRT-1 administered exogenously had no additional effect in the modulation of maternal behavior.

Recently, Diniz et al. (2018) showed a greater number of HCRT-immunoreactive neurons in lactating dams compared to that of virgin females, and a decrease from PPD15 to PPD21 in response to regular suckling stimulus of pups, suggesting that a role of HCRTergic system during the lactation period (Diniz et al., 2018). However, the specific role that HCRT play in maternal behavior remains to be elucidated.

### HCRT, mPOA and thermoregulation

The higher dose of HCRT-1 (HCRT_200_) increased body temperature during the first and second hour after its microinjection into mPOA. This is in accordance with elevated body temperature after infusion of HCRT-1 intracerebroventricular in rats, with a peak at about 3 hours (Yoshimichi et al., 2001). Regarding the POA, only one study has administered HCRT-1 (1mM) into lateral preoptic area of male rats, but brain temperature was not affected by HCRT-1 (Methippara et al., 2000). These data, together with the present results, suggest that the HCRT modulates body temperature acting specifically within the mPOA rather than in other POA areas, supporting the large body of evidence that establish the mPOA as a key center of body temperature regulation (Boulant, 2000; Harding et al., 2018; Morrison, 2016). However, blocking endogenous HCRT action by the administration of DORA had no effect on body temperature of the mothers. This latter result could suggest that endogenous HCRT may not be playing a main role in maintaining basal temperature throughout the mPOA circuit in lactating dams, which could reflect the changes in the mPOA circuits during motherhood and lactation. In this sense, it is important to note that functional differences in mPOA networks have been reported between male and postpartum female rats (Bleier et al., 1982; Brown et al., 1988; Gorski et al., 1978; Ottem et al., 2004; Raisman and Field, 1971).

In the present report, the increase in maternal body temperature provoked by HCRT_200_ was not accompanied by changes in the time spent in nursing. Although (Leon et al., 1978) suggest that nursing bouts are limited by a rise in maternal temperature, and that the rate of temperature rise determines the duration of each bout, other evidence show that hyperthermic mothers (following treatment with morphine plus naloxone) nurse their litter normally. These results suggest that nursing bouts in lactating rats are not limited by the mother’s temperature (Stern and Azzara, 2002). Thus, body temperature and nursing integration are still to be determined.

## Conclusions

Our work shows that in lactating rats, HCRT-1 modulates mPOA to promote wakefulness, whereas DORA promotes both NREM and REM sleep. These effects of DORA on sleep parameters are accompanied by an increase in the time that mother rats nurse their litter and in the number of milk ejections. HCRT-1 into mPOA promoted wakefulness that was associated with a slight increase in body temperature, whereas DORA did not alter body temperature in lactating mother rats. Taken together, our results suggest that the reduction of the endogenous HCRT within the mPOA of lactating rats is important to promote sleep, nursing and milk ejection simultaneously, without affecting body temperature, while external additional HCRT mainly promotes wakefulness and increase body temperature in this behavioral state. Overall, this study highlights the role of the hypocretinergic system, by acting through mPOA neurons, in the modulation of sleep, maternal behavior and body temperature in an integrated fashion.

## Acknowledgements

This work was partially supported by “Programa de Desarrollo de Ciencias Básicas (PEDECIBA)” and “Agencia Nacional de Investigación e Innovación (ANII)”. All authors have seen and approved the manuscript, and it hasn’t been accepted or published elsewhere. The authors have no competing interests.

